# Glutamatergic projections from ventral hippocampus to nucleus accumbens cholinergic neurons are altered in a mouse model of depression

**DOI:** 10.1101/2022.04.20.488950

**Authors:** Lucian Medrihan, Margarete Knudsen, Tatiana Ferraro, Pedro Del Cioppo Vasques, Yevgeniy Romin, Sho Fujisawa, Paul Greengard, Ana Milosevic

## Abstract

The cholinergic interneurons (ChATs) of the nucleus accumbens (NAc) have a critical role in the activity of this region, specifically in the context of major depressive disorder. To understand the circuitry regulating this behavior we sought to determine the areas that directly project to these cells/interneurons by utilizing the monosynaptic cell-specific tracing technique. Mapping showed monosynaptic projections that are exclusive to NAc ChATs. To determine if some of these projections are altered in a depression mouse model, we used mice that do not express the calcium binding protein p11 specifically in ChATs (ChAT-p11 cKO) and display a depressive-like phenotype. Our data demonstrated that while the overall projection areas remain similar between wild type and in ChAT-p11 cKO mice, the number of projections coming from the ventral hippocampus (vHIP) is significantly reduced in the ChAT-p11 cKO mice. Furthermore, using optogenetics and electrophysiology we showed that glutamatergic projections from vHIP to NAc ChATs are severely altered in mutant mice. These results show that specific alterations in the circuitry of the accumbal ChAT interneurons could play an important role in the regulation of depressive-like behavior, reward seeking behavior in addictions, or psychiatric symptoms in neurodegenerative diseases.

## Introduction

Nucleus accumbens (NAc) is a large nucleus at the ventral part of the striatum, where signals from a wide variety of sensory and motor inputs converge. The NAc functions as a funnel through which information must pass and where it is finely modulated ^1, 2^. The NAc circuitry is centrally involved in the assessment of reward. It selects and reinforces the behaviors that lead to a positive outcome and abandons or modifies those that result in non-productive activities, and thus has a major role in generation of motivated behaviors ^3, 4^. Disrupted functioning of this network has been linked to the incapacity of experiencing pleasure (anhedonia) as seen for example in depression ^5, 6^. Thus, mapping complex NAc neuronal networks with other brain regions would be fundamental for understanding the behavioral changes in mood disorders and other neurological diseases. NAc connectivity was examined using the morphological and electrophysiological approaches, as well as non-selective anterograde and retrograde tracers ^7-11^. These classical studies showed that the main brain areas that project to the NAc are the prefrontal cortex (PFC), ventral hippocampal subiculum (vHIP/Sub), basolateral amygdala (BLA), ventral tegmental area (VTA), and thalamic nuclei ^12, 13^. These areas reciprocally interact through excitatory glutamatergic inputs of the cortico-striato-pallidal-thalamo-cortical loop, and by the dopaminergic projections from the ventral tegmental area (VTA) and substantia nigra (SN) in the mesolimbic dopaminergic pathway. Although the connections of the NAc have been well characterized, the direct cell-cell interaction with other brain areas is still not well understood. Studies showed that a number of/numerous diseases, major depressive disorder among them, have a strong cell-specific and region-specific component ^14^, making it necessary to decipher the cell type specific circuitry between the key brain regions. The gross anatomy of inputs to numerous brain areas has been studied using traditional tracers ^11, 15, 16^, but these techniques cannot distinguish connectivity to specific cell types. This has become achievable with the advent of techniques that allow the cell-specific connectivity mapping ^17^, enabling the study of the processes in only one cell type without dissecting it out from its natural surroundings, preserving all the connections and processes that exist in vivo. Monosynaptic circuit tracing with the CRE-dependent modified rabies virus, developed by the Callaway laboratory ^18^ and modified by the Uchida laboratory ^19^, circumvent this limitation by using mouse lines that express CRE in a cell-specific manner. With the introduction of this cell-based approach for mapping connectivity inputs to different cell types in the dorsal striatum, including D1 and D2 medium spiny neurons, ^20^, and ChAT interneurons ^21, 22^ were elucidated. However, the cell-specific inputs to the ventral striatum have not been studied.

Striatal cholinergic (ChAT) neurons are giant aspiny interneurons that represent only about 0.3% of all striatal neurons ^23^. Despite their under-representation, the tonically active ChAT interneurons have a dominant modulating control over inputs and outputs of the main population of neurons in NAc, the medium spiny neurons (MSNs) ^5, 24^. Indeed, optogenetical inhibition of ChAT interneurons in the NAc increased MSN activity and blocked the rewarding effects of cocaine ^25^. Chemogenetic and optogenetic manipulation of NAc ChAT interneuron activity opposed the motivating influence of appetitive cues, further emphasizing their importance for cue-motivated behaviors ^26^. These cells have been also shown to have a critical role in regulating depressive-like behaviors in mice ^27-29^. One of the molecular mechanisms of depressive-like behaviors in mice are the alterations in expression levels of the calcium-binding protein S100a10 (also called p11), shown by our laboratory and others ^30, 31^. Mice with a complete lack of p11 exhibit anhedonia and despair, but this depressive-like behavior can be reiterated with the region-specific knock down in NAc ^27^, and specifically in the NAc ChATs ^14^. Notably, this behavior appears to be governed specifically by the NAc and not CPU ChATs, since selective p11 knock down in CPU ChATs did not reproduce the depressive phenotype.

For all these reasons, it is important to examine the connectivity of these neurons in order to gain a better understanding of cell-specific circuitries regulating normal and pathologic behavioral outcomes. Here, we utilized the CRE-dependent monosynaptic modified rabies virus method and mapped all the brain regions and cells with direct inputs to NAc ChATs. We corroborated known areas with direct inputs to NAC, but also found areas that project only to NAc but not the caudate putamen-dorsal striatum (CPU) ChATs. To trace the direct inputs to ChATs in the context of depression, we used ChAT-CRE mice with the depressive phenotype (ChAT-p11 cKO). We found that in ChAT-p11 cKO mice a significantly lower number of pyramidal cells from the ventral hippocampus projects to the NAc ChATs. Together these data demonstrate the inputs to ChAT neurons from a large number of discrete areas scattered throughout the brain and provide possible mechanism of how accumbal ChAT circuitry regulate depression and other NAc-related behaviors.

## Results

We sought to determine direct inputs onto the ChAT neurons of the nucleus accumbens using the monosynaptic circuit tracing method with the CRE-dependent modified rabies virus technique ^17, 19^. This method utilizes the spread of rabies virus (RV) through the synapses onto the projecting neurons. However, this spread is limited to the direct, primary input onto the infected cells due to several modifications in the rabies virus genome. We used the previously characterized ChAT-CRE mouse line GM60 (www.gensat.org). Experimental design was adapted to allow the optimal infection of the sparce cell population. ChAT cells were labeled with stereotaxic injection of a mixture of two viruses, one carrying the G-protein and another carrying the TVA receptor (UNC Gene Therapy Center, Chapel Hill, NC). This was followed by injection of a rabies virus into the same NAc region five weeks later (Fig. 1A and B). We confirmed that helper viruses, carrying the mCherry tag, infected ChAT cells by immunocytochemical labeling with anti-mCherry and anti-ChAT antibodies (Fig. 1C). Next, we confirmed that viral injections infected only NAc ChAT cells using the mouse line ChAT-CRE crossed to ChAT-TRAP (www.gensat.org) in which all ChAT cells are tagged with GFP ^33^. We immunolabeled series of coronal sections with an anti-mCherry antibody and detected no mCherry+ GFP+ cells in the neighboring brain regions with ChAT cells (Fig. 1D). Furthermore, because the injection into NAc infects the dorsal striatum-caudate putamen (CPU) ChAT+ cells along the needle track, a set of control injections into the CPU were performed to determine the overlap with the CPU ChATs projections.

**Figure 1.**
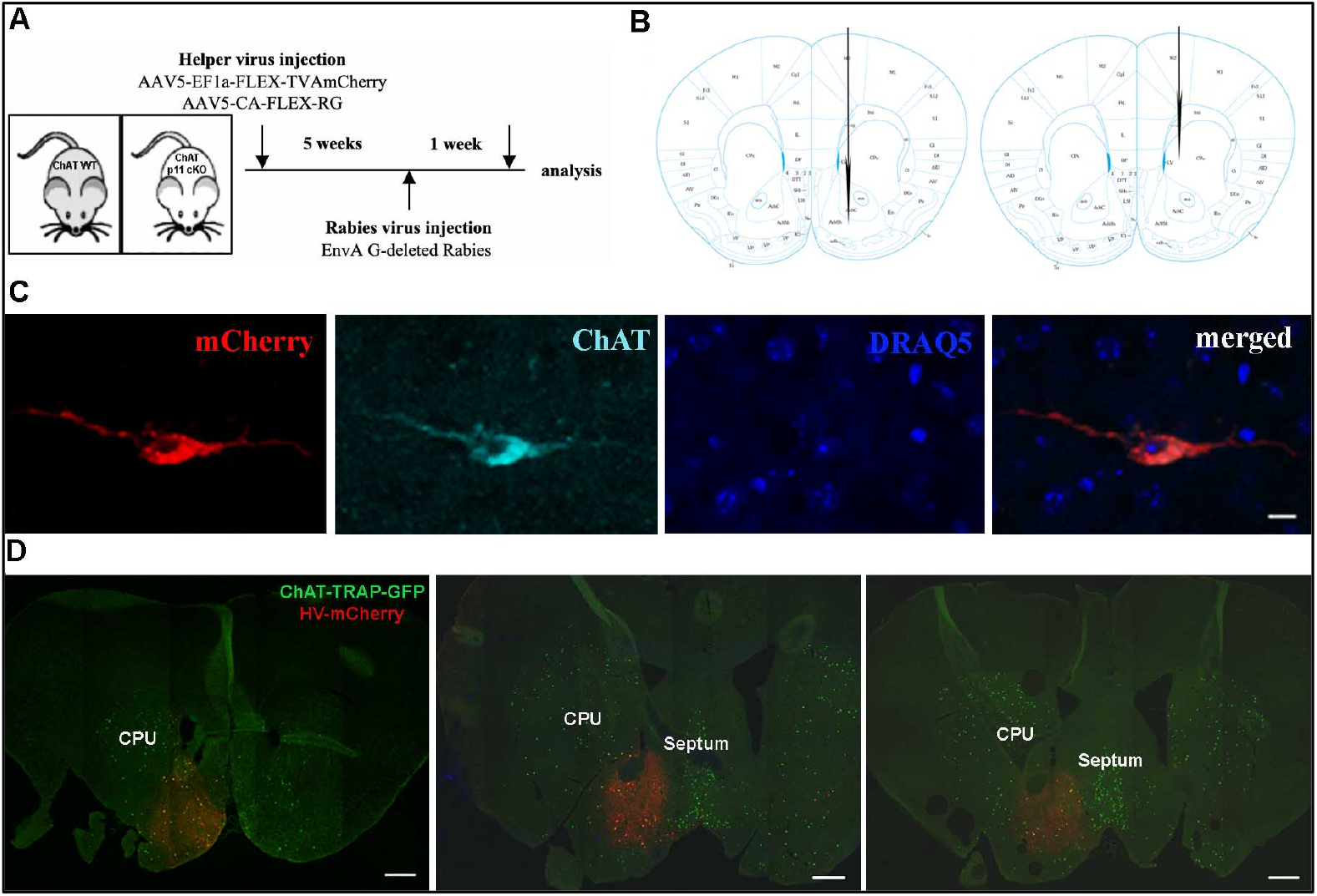
ChAT cells in NAc are successfully infected utilizing the rabies monosynaptic tracing method: **A)** Injection schedule and types of viruses used for axonal tracing. **B)** Schematic diagram showing the areas of the brain injected with the viruses. **C)** Representative image of the cell in the NAc infected with the mixture of helper viruses (red, mCherry) and immunolabeled with the anti-ChAT antibody. Scale bar 10 µm. **D)** Series of low magnification images of the sagittal sections of the mouse line ChAT-CRE crossed with the ChAT-L10a-EGFP injected with the helper (mCherry) virus into the NAc. Sections were labeled with the mCherry (red) and GFP (green) antibodies to enable visualization of the infection efficacy in the ChAT cells. Majority of infected cells, double labeled with mCherry and GFP (yellow) are found in the NAc, with no major infection of the ChATs in the adjacent areas of dorsal striatum, basal forebrain, and septum. Scale bars are 500 µm.

To generate a comprehensive list of all projection sites and to quantify the number of neurons that project to NAc ChATs we analyzed the whole brain, as described in detail in the methods section. The cells retrogradely labeled with RV-GFP virus were detected in distinctive subregions of the cortex, hippocampus, thalamus, hypothalamus, amygdala, and brainstem (Fig. 2, Suppl. Fig 1A). The representative images of the NAc ChAT projection sites are shown in Fig. 2A-G, while a complete list of all regions and proportions of labeled cells in each area are shown in Fig. 2H. In addition, to gain the insight into the similarities and differences between the NAc and CPU projections, we also analyzed the brains injected into the CPU (Suppl. Fig. 1B). Notably, the regional distribution of cells labeled with the rabies virus after the CPU and NAc injections had some very distinct areas with labeled cells (Suppl. Fig 1A, B). This suggested that NAc ChATs have distinct set of regions that project exclusively to these cells and not the CPU ChATs.

**Supplemental Figure 1.**
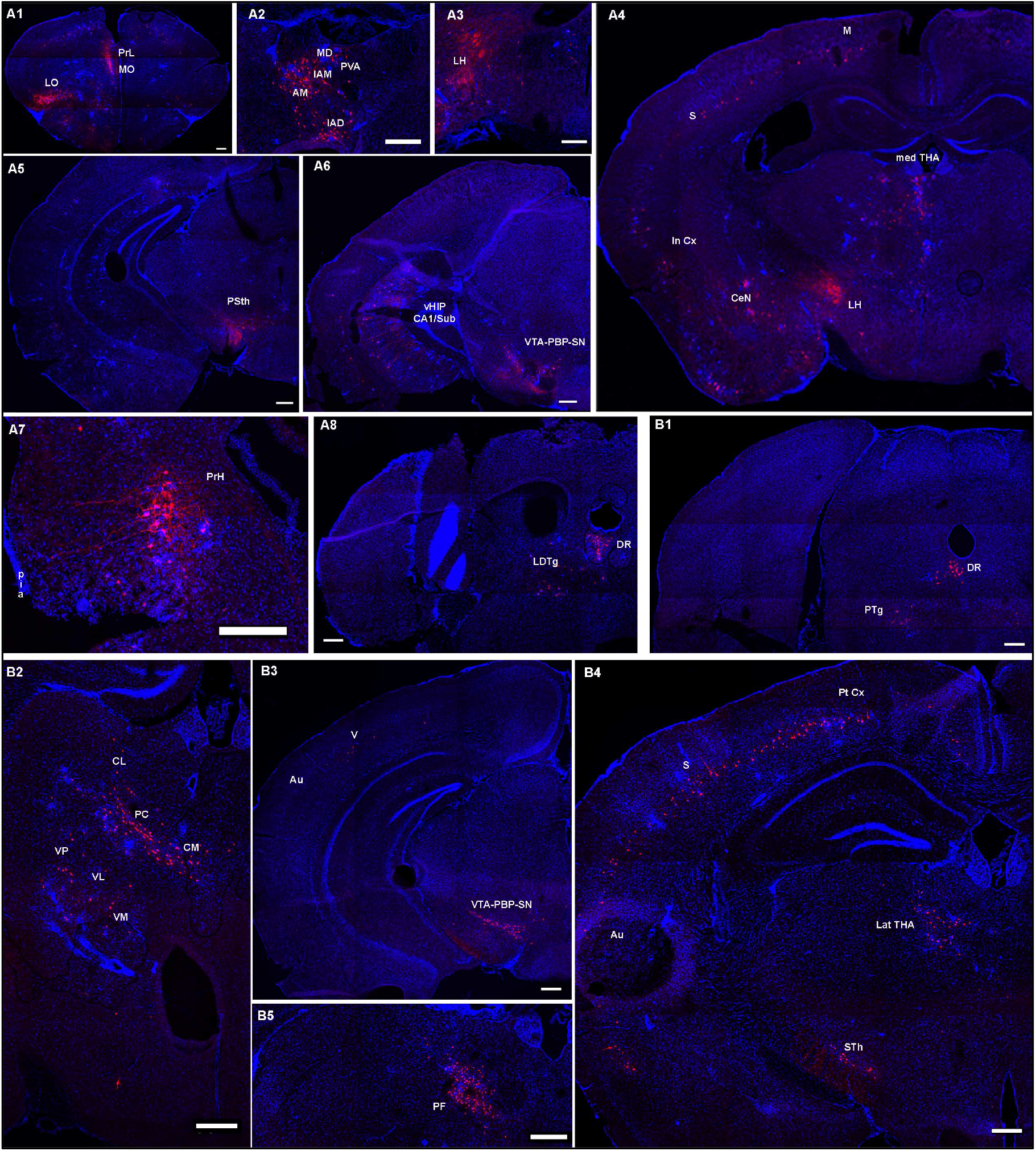
Monosynaptic inputs to ChAT cells in the ventral and dorsal striatum show regional differences. Representative areas from the coronal sections of the brain injected into the NAc ChATs **(A1-8)** and CPU ChATs **(B1-5)** with the RV-mCherry. **A1)** The whole brain section of the prefrontal cortex. **A2)** Projecting neurons in thalamic nuclei PV, CM, MD, PC, IAD, IAM. **A3)** Lateral hypothalamic area with large number of projecting neurons. **A4)** The whole brain section showing projecting neurons in the cortical areas M1 and 2, S1 and S2, insular cortical areas, central amygdaloid nuclei, ZI, and thalamic and hypothalamic nuclei. **A5)** Area of the parasubthalamic nucleus contains projecting cells, while in the hippocampus small number of cells can be observed in the subiculum and CA1 area. **A6)** In the mor caudal area of the ventral hippocampus large number of labeled cells are found in the subiculum – CA1 regions. Cells are also visible in the VTA-PBP area. **A7)** Perirhinal cortical area contains large number of pyramidal neurons labeled with mCherry. **A8)** Tegmental nuclei and dorsal raphe contain labeled cells in the lateral tegmental nuclei, and parabrachial nuclei. **B1)** Labeled cells in the tegmental area that project to the CPU ChATs. Notice absence of the labeled cells in the subiculum of the ventral hippocampus. **B2, B5)** Thalamic regions of the CPU injected brain depicts different thalamic nuclei that project to the CPU ChATs. **B3)** Substantia nigra, VTA-PBP-SN area contains large number of cells labeled with the mCherry. Note the absence of labeled cells in the hippocampus. **B4)** Whole brain image showing large number of cortical areas that contain labeled cells, and strong labeling in the subthalamic and parasubthalamic nuclei. Scale bars in all images are 100 µm.

**Figure 2.**
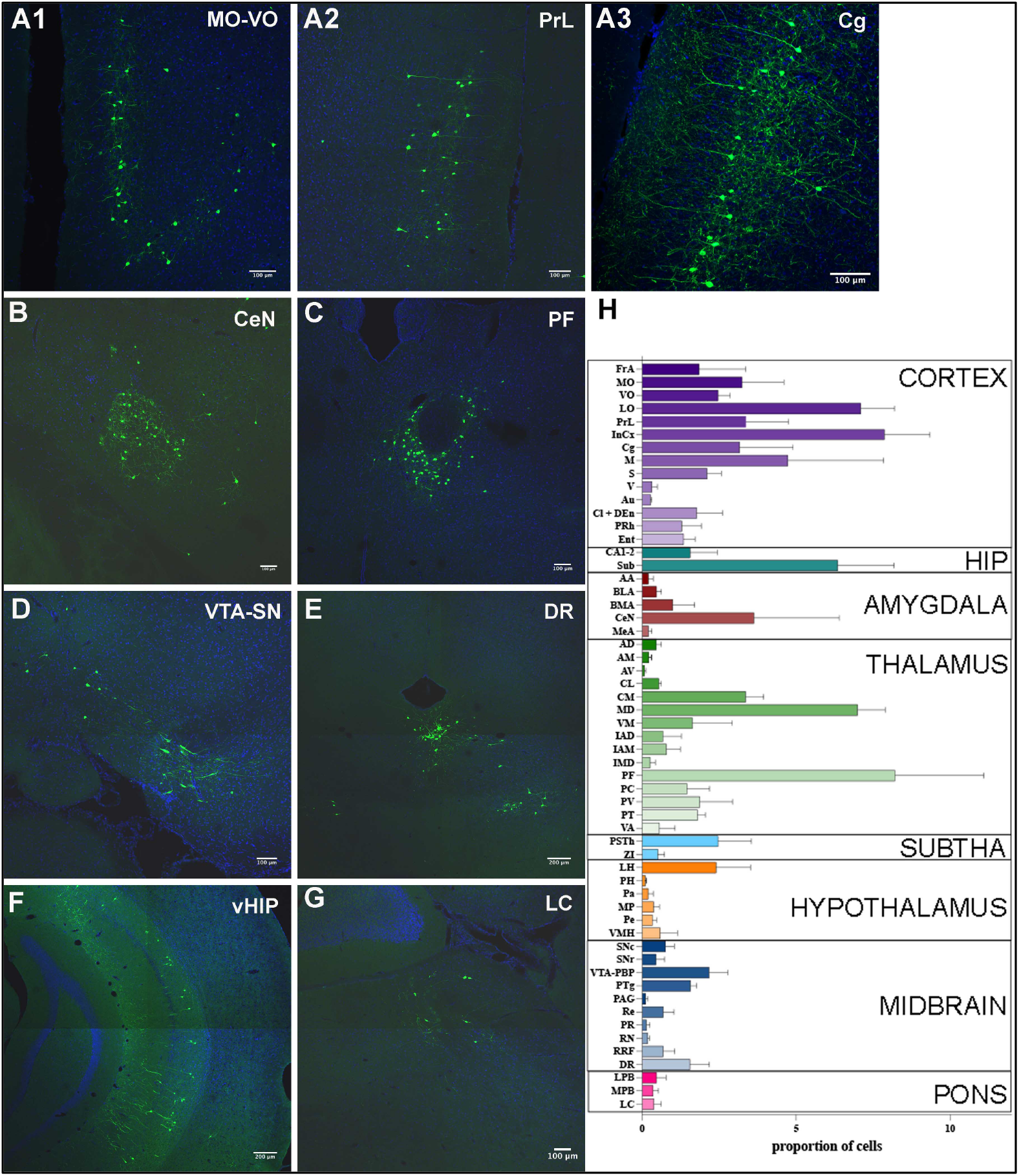
Representative images of the brain areas that project directly to the NAc ChATs and that mediate depressive-like phenotype. Anti-GFP antibody immunolabeling revealed cells in several cortical areas including medial (MO) and ventral (VO) orbital prefrontal areas shown in **A1**, prelimbic cortex (PrL) in **A2** and cingulate cortex (Cg) in **A3**; **B)** central amygdaloid nucleus (CeN), **C)** parafascicular thalamic nucleus (PF), **D)** ventral tegmental area and substantia nigra (VTA-SN), **E)** dorsal raphe (DR), **F)** ventral hippocampus (vHIP), and **G)** locus coeruleus (LC). **H)** Graph represents the proportion of RV-GFP infected cells quantified throughout the brain of mice injected in the NAc. Graph bars are represented as data mean ± SEM from three replicate samples. HIP - hippocampus, SUBTHA – subthalamus.

### Cortical projections to NAc ChATs

Large proportion of labeled cells in the brains that have been injected into NAc was found in the discrete areas of the prefrontal cortex (Fig. 2H). These included medial (Fig. 2A1) and lateral orbital cortex (Suppl. Fig 1A1), agranular insular cortex, prelimbic cortex (Fig. 2A2) and anterior cingulate cortex (Fig. 2A3). Majority of the projections are coming from the ipsilateral side (Suppl Fig. 1A), as opposed to CPU where a substantial projection from the contralateral side was reported ^36^.

In the orbital areas of the prefrontal cortex NAc ChAT projections were in the MO-VO, while they were absent in the CPU ChAT projections (Fig. 3A1-2). Prefrontal cortical areas contained large number of labeled neurons in the layer 2/3 (Fig. 2A1-2, Fig. 3A1), opposite to the labeled cells after CPU injection that concentrated in the deep cortical layers (Fig. 3A2 and ^21, 22^. To confirm that layer 2/3 cells in MO-VO were indeed glutamatergic pyramidal cells that project to NAc ChATs two sets of experiments were performed. Firstly, we adapted the injection approach to test the connectivity. For this purpose, we used another mouse line with a CRE expression under the regulation of the Cux2, specific marker for layer 2/3 pyramidal neurons ^37^. Helper virus, tagged with mCherry, was injected into the NAc of the Cux2-CRE mouse line (gift from Dr. Muller, Johns Hopkins, Baltimore, MD), followed by the injection of the RV-GFP into the upper layers of the MO-VO five weeks later, as shown in the schematic diagram in Fig. 3B. This injection approach enabled layer 2/3 projecting neurons to be infected with the helper virus via axonal ends in the NAc. The consecutive RV-GFP injection and recombination enabled projecting neurons labeled with the helper virus to express the GFP. The resulting double labeled cells (mCherry/GFP) in the medial orbital cortex layer 2/3 are shown in Figure 3C. Secondly, a set of electrophysiological recordings were done to confirm our initial data (Fig. 3D-H). We used whole-cell patch clamp to record from layer 2/3 neurons infected with RV-GFP and helper virus, respectively. The firing frequency, as well as the membrane potential and action potential threshold of the recorded neurons were characteristic to pyramidal neurons (Fig. 3D) and were not changed between neurons infected with the different viruses (Fig 3E-H). These experiments confirmed that layer 2/3 in the MO-VO are indeed pyramidal projecting neurons and that their basic physiological properties are not altered by the virus injection protocol. Interestingly, Wfs1-expressing cells with the same layer and regional specificity project to the motor cortex and CPU, and receive projections from the lateral amygdala, posterior thalamic group, and several cortical areas ^38^, suggesting that multiple subtypes of the layer 2/3 neurons from the medial orbital cortex project outside of the cortex.

**Figure 3.**
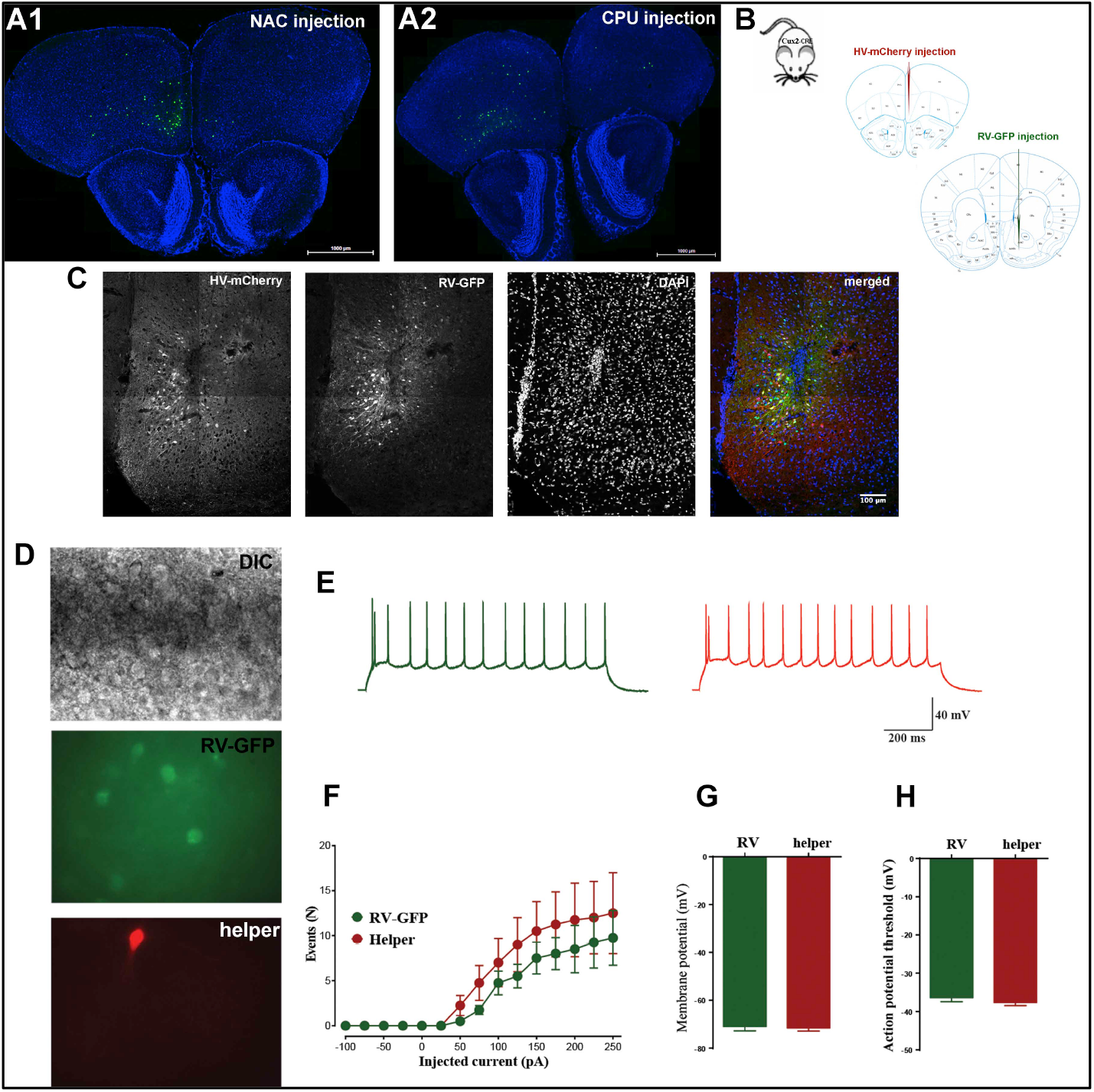
Layer 2/3 pyramidal neurons from the prefrontal cortical areas project to NAc ChATs. **A1-A2)** Low-magnification images from the prefrontal cortex MO-VO showing layer 2/3 neurons **(A1)** in contrast to layer 5 neurons that project to CPU ChATs **(A2). B)** Experimental design of injections into the Cux2-CRE mouse line to label the medial orbital cortex MO layer 2/3 neurons that project to NAc ChATs. Mouse was first injected with the helper virus (mCherry) into the MO, followed by injection of RV-GFP into the NAc. **C)** Immunocytochemistry of the MO cortex of the Camk2a-CRE mouse labeled the anti-mCherry and anti-GFP antibodies, showing that some cells infected with the helper virus (mCherry+) and rabies virus (GFP+) are double labeled (mCherry+ GFP+, merged). **D)** Representative cells in the layer 2/3 of the CaMK2a-CRE mouse prefrontal cortex labeled with the helper virus into the MO-VO followed by the RV-GFP injection into the NAc that were used for electrophysiology recordings. **E)** Representative cells with traces of the firing response to a 250-pA step injection of current in layer 2/3 MO-VO pyramidal neurons infected with RV-GFP (GFP, green) and helper virus (mCherry, red). **F)** The representative firing traces firing frequency of layer 2/3 MO-VO pyramidal neurons infected with RV-GFP (*n* = 4neurons/3mice) or helper virus (4neurons/3mice) in response to 50-pA current steps injections. **G-H)** Histograms showing the membrane potential (F) and the voltage threshold (H) for action potential firing in layer 2/3 MO-VO pyramidal neurons infected with RV-GFP or helper virus.

### ChAT circuitry is changed in a depressive-like mouse model

To better understand the circuitry at the root of depressive behavior we investigated projections between control and depressive-like mouse model ChAT-p11 cKO mice. Quantitative analysis was performed on four brain regions: MO-VO, AMY, vHIP, and DR. These areas were selected because they have been shown to have an important role in depressive-like behaviors ^6, 39-41^, and because MO-VO, AMY and vHIP project specifically to the NAc ChATs. Representative images of projections from MO-VO, vHIP and DR used for this quantification are shown in Fig. 4A. Quantification of projections have shown a significantly larger number of projections from vHIP in the ChAT WT mice that in Chat-p11 cKO (Fig. 4B). A significant difference was not reached for MO-VO, AMY, and DR, although there is a trend towards lower number of projections from AMY, and higher number of projections from MO-VO in ChAT-p11 cKO.

**Figure 4.**
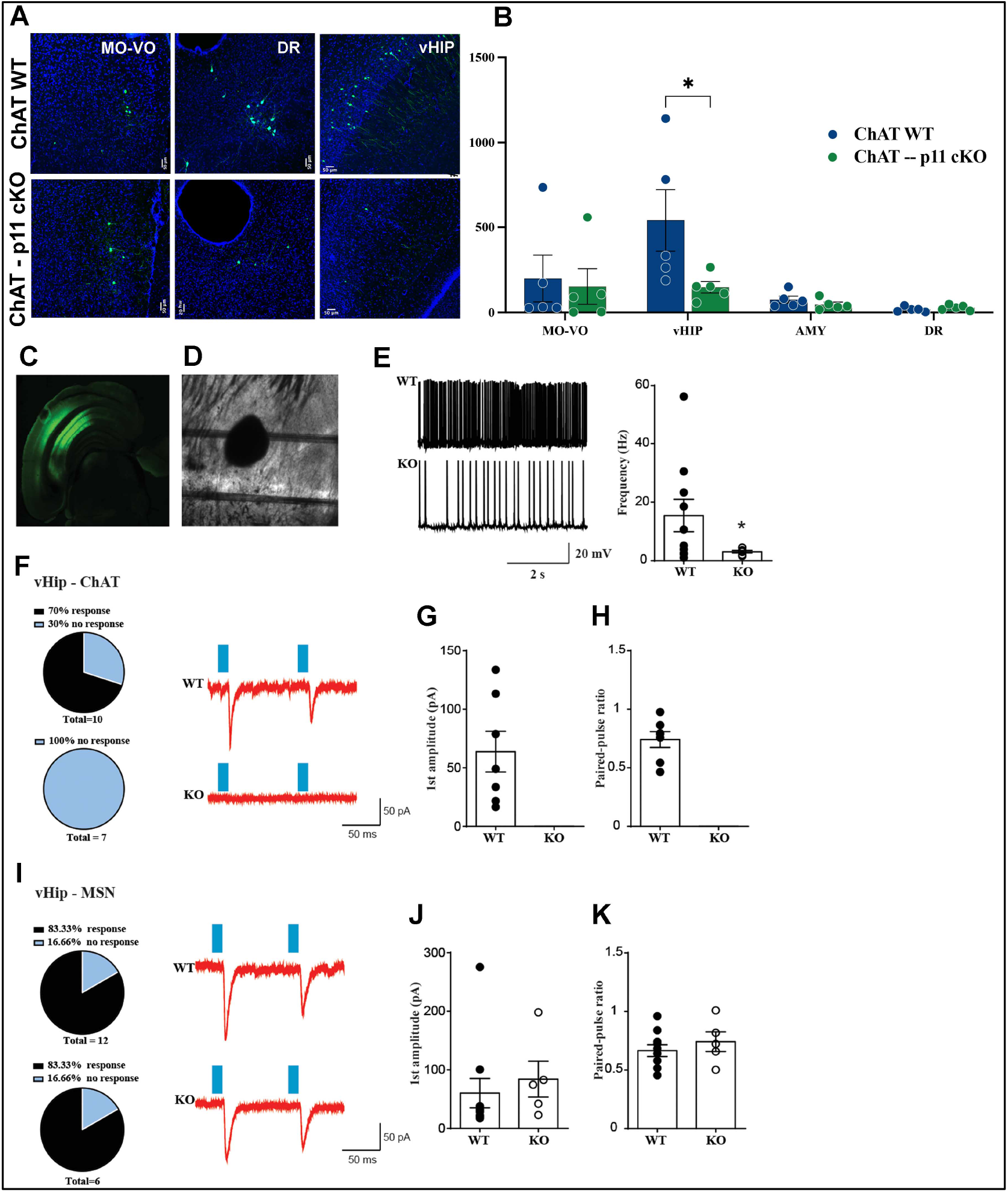
Mouse model of depression – ChAT-p11 cKO shows differential projections to NAc ChATs: **A)** Representative images of projection neurons labeled with RV-GFP in MO-VO region of the prefrontal cortex, DR, and vHIP in ChAT-CRE and ChAT-p11 cKO mice lines. The injections were designed to infect small number of cells to allow optimal quantification of projecting neurons. **B)** Quantification of projections from MO-VO, DR, AMY and vHIP regions. **C)** Low-magnification image of ventral hippocampus showing CaMK2a-ChR2 (H134R) –eYFP infection CA1 and subiculum pyramidal neurons. **D)** Differential interference contrast image showing the area from NAc where we recorded from ChAT and MSN neurons. **E)** Representative traces of tonic firing of ChAT neurons from WT and p11 KO mice and histograms quantifying the firing frequency of recorded neurons from each genotype, * p< 0.05, Kolmogorov-Smirnov test (*n* = 10 neurons/4 mice for WT and 8/3 for p11 KO). **F)** Pie chart showings the percentage of WT and p11KO ChAT neurons that responded to ChR2 stimulation of vHIP terminals in NAc and representative paired-pulse (100 ms interstimulus) traces in response to ChR2 field light stimulation. **G)** Histograms showing the amplitude of the first response (gE) and the paired-pulse ratio at 100 ms interstimulus **(H)** in response to ChR2 field light stimulation of NAc in WT and p11KO ChAT neurons. **I)** Pie chart showings the percentage of WT and p11KO MSN neurons that responded to ChR2 stimulation of vHIP terminals in NAc and representative paired-pulse (100 ms interstimulus) traces in response to ChR2 field light stimulation. **J-K)** Histograms showing the amplitude of the first response (**J**) and the paired-pulse ratio at 100 ms interstimulus (**K**) in response to ChR2 field light stimulation of NAc in WT and p11KO MSN neurons.

Virtually all labeled cells in the hippocampus were confined to the ventral part, namely subiculum and CA1. No direct input to CPU ChATs was detected. This corroborated and expanded on previous findings that ventral subiculum has a strong projection to NAc ^7, 8, 10^. To verify if the connections from vHIP to the NAc ChAT neurons are functionally altered by the p11 deletion we expressed channelrhodopsin (ChR) in ventral hippocampus pyramidal neurons (Fig 4C, D) of the Camk2a-CRE mouse. We have shown before that deletion of p11 from ChAT neurons results in a decrease in their firing frequency ^28^, so we used this to identify the ChAT neurons by their tonic firing in acute slices containing NAc from WT and p11 KO mice (Fig. 4E). Next, we recorded photostimulation-evoked glutamatergic currents of the vHIP terminals in NAc ChAT neurons from WT and p11 KO mice. We applied a paired-pulse protocol with two light pulses (1-2 ms) separated by 100 ms interval to verify first if the connections between vHIP pyramidal neurons and NAc neurons are functional and second if the connections show normal short-term synaptic plasticity. While in the WT ChAT neurons we were able to elicit glutamatergic postsynaptic responses in 70 % of the recorded neurons, we could not find any response in p11 KO ChAT neurons in multiple mice (Fig. 4F-H). To validate our recording protocol, we took advantage of the well-established connection between medium spiny neurons (MSN) and HIP ^8, 10^ by recording photostimulation-evoked glutamatergic currents in neighboring MSN neurons from the same slices. Using this paired-pulse protocol we evoked glutamatergic responses in 83.3% of the tested MSN neurons in both WT and p11 KO mice and both the amplitude of the response and the paired-pulse ration was unchanged between the genotypes (Fig. 4I-K). Taken together, these data show for the first time the existence of functional glutamatergic synaptic connections between vHIP pyramidal neurons and further emphasize the alterations of vHIP ChAT projections induced by the deletion of p11 in ChAT neurons.

### Accumbal ChATs receive inputs from many brain areas regulating complex behaviors

Large number of areas contained cells labeled by retrogradely transferred rabies virus after injections into the NAc and CPU (Suppl. Fig 1A, B).

Labeled cells were distributed in distinct areas of the **amygdala**. In NAc injected brains central amygdaloid nucleus prominently displayed large number of GFP+ cells (Fig. 2B, Suppl. Fig 1A). When brains injected into the NAc and DStr were examined labeled cells were distributed in distinct areas. In NAc injected brains central amygdaloid nucleus (CeN) contained vast majority of labeled cells (Fig. 2B). We observed very small number of GFP+ cells in the BLA, suggesting that these inputs synapse on other types of accumbal neurons. It is well-established that amygdala, especially basolateral amygdaloid nucleus project to the striatum ^39^. More recently, cell-based tracing methods showed that both central and basolateral amygdala project specifically to Drd1 MSN in the dorsal striatum ^20^. Functional studies investigating role of amygdalar subdivisions linked BLA and CeN to fear conditioning ^42, 43^ and memory consolidation ^44^, while lesions of the CeN reduced stress response and anxiety and fear response to chronic unpredictable stress ^45^. In the brains injected into the DStr labeled neurons were predominantly in the anterior cortical, central, and medial amygdaloid nuclei. Interestingly, we did not observe cells in basolateral or central amygdaloid nuclei, suggesting distinct inputs to ChAT neurons from dorsal and ventral striatum. The data presented here demonstrating strong connectivity of central amygdaloid nucleus to the NAc ChATs, and connectivity of basolateral amygdala to other accumbal cells would suggest the intricate modulation of fear conditioning, anxiety, and stress responses in striatum.

Majority of labeled cells in the **thalamus** of the brains injected into the NAc was concentrated in parafascicular, mediodorsal and centromedial thalamic nuclei (Fig. 2C, Suppl. Fig. 1A2, A4). It is notable that inputs to the CPU ChATs are coming from a different set of nuclei (Suppl. Fig. 1B2, B4, B5, and ^21, 22^), with few exceptions such as parafascicular nucleus that projects to both dorsal and ventral striatal ChATs. Others have shown that both D1R and D2R MSN receive afferents predominantly from parafascicular and mediodorsal nuclei, as well as smaller number of inputs from ventromedial, anterodorsal and anteroventral nuclei ^20^. This would imply that thalamo-striatal connectivity target both projection neurons and interneurons, bringing additional level of complexity in control over striatal activity.

**Hypothalamus** of mice injected in the NAc contained very small number of labeled cells, scattered mostly in lateral hypothalamic area (Fig. 2H, Suppl. Fig. 1A3). Lateral hypothalamic area has been shown to connect with basomedial and anterior cortical amygdaloid nuclei ^46^ and has a role in food intake regulation ^47, 48^. This would suggest that inputs to ChAT neurons in NAc have a role in modulating the behavior related to feeding and/or food intake.

Strong labeling of neurons in **substantia nigra (SN) and ventral tegmental area (VTA)** after NAc injection were observed (Suppl. Fig 1A6). The striatal-VTA circuitry was implicated in the regulation of stimulus dependent learning ^49^, as well as reward behavior ^50^. The projections from SN and VTA to the neurons of the dorsal striatum cells are well documented ^20-22^. Here we have shown for the first time that the part of this circuitry involves the SN/VTA-NAc ChAT connectivity.

**Dorsal raphe** contained labeled cells in the brains from NAc and CPU injection. This affirmed previous findings about relatively weak serotoninergic inputs into striatum ^51^, and specifically ChAT cells ^21, 22, 52^. Interestingly and not previously reported, we found virally labeled cells to be confined only to the ipsilateral side of the brain (Fig. 2E).

Medulla, and specifically locus coeruleus, the main noradrenergic output in the brain contained few labeled cells. In addition, we detected very few neurons in other areas of pons and medulla, most notably area of the gigantocellular reticular nucleus. However, these neurons were sparse and thus difficult to be assigned to the appropriate nuclei. This is probably due to the technical limitations of the technique used in this study but may also represent the actual *in vivo* representation of connectivity within the brain.

## Discussion

Mapping complex networks between the neurons in different regions of the central nervous system is essential to understand the behavior and its changes in neurological diseases. Here, we have generated a detailed and accurate/exhaustive brain-wide map of direct projections to the cholinergic neurons in the nucleus accumbens (Fig. 5A). Importantly, this mapping revealed several areas that project only to the ventral but not dorsal striatal ChATs, such as layer 2/3 from the medial and ventral subdivisions of the orbital prefrontal cortex, central amygdaloid nucleus and subiculum of the ventral hippocampus. Most importantly, the circuitry between NAc ChATs and ventral hippocampus is significantly altered in the mouse model of depression, ChAT-p11 cKO, with implications for cognizing the specific morphological and anatomical changes in psychiatric diseases involving these brain networks.

**Figure 5.**
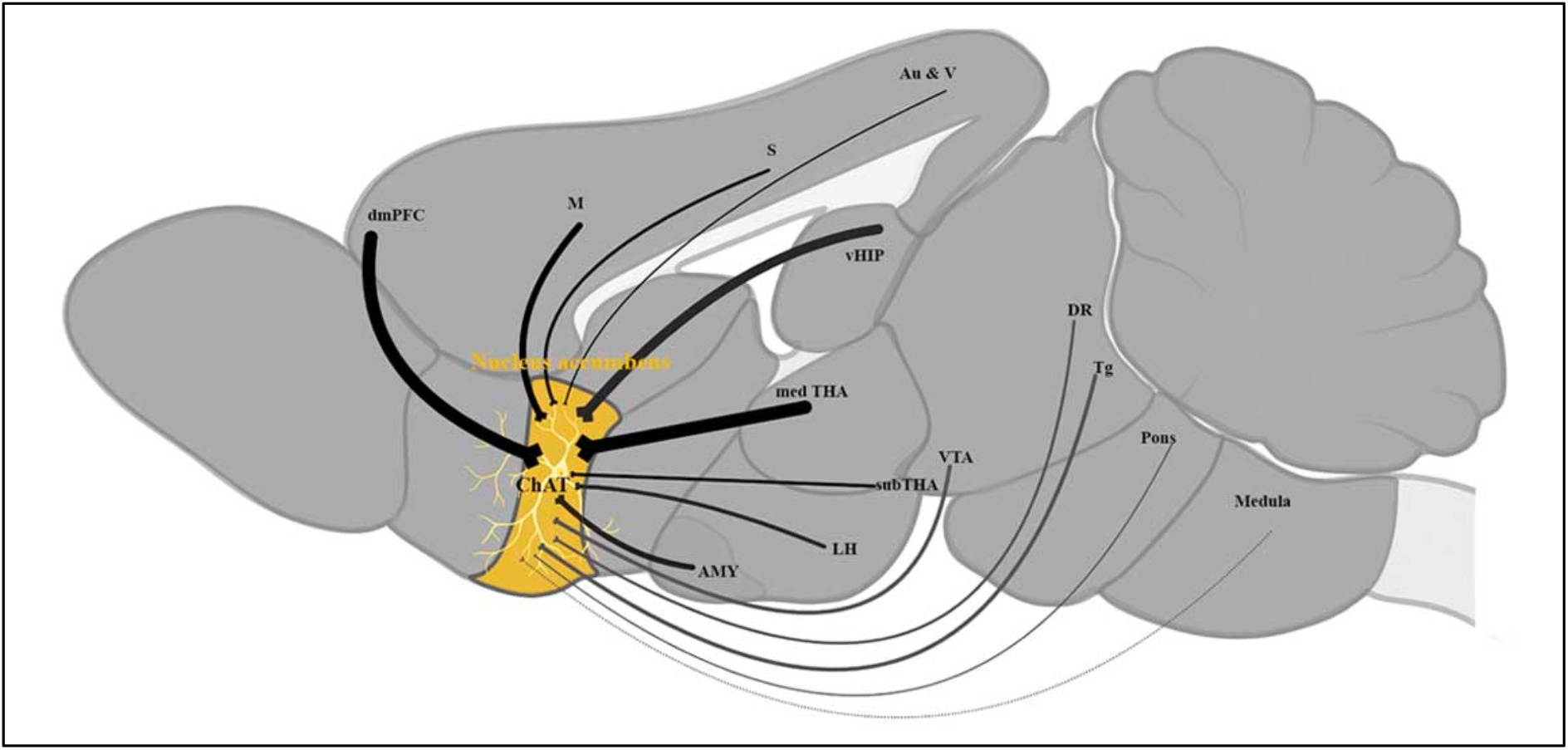
ChATs of the nucleus accumbens receive large number of projections from multiple regions of the brain. Summary of the major projections to the ChAT cells of the nucleus accumbens. The line width represents the proportion of cells in that brain region projecting to the nucleus accumbens, relative to other regions. dmPFC – dorsomedial prefrontal cortex; M – motor cortex; S – somatosensory cortex; Au & V – auditory and visual cortices; vHIP – ventral hippocampus; medTHA - medial thalamic nuclei; LH - lateral hypothalamus, AMY – amygdaloid nuclei; subTHA - subthalamic region; VTA – ventral tegmental area; DR - dorsal raphe; Tg – tegmental area.

The NAc is involved in mediating an altered perception of social status ^53^, inability to experience pleasure and positive emotions, heighten negative emotional processing, psychomotor retardation ^54^, and behaviors related to reward ^5^, all distinctive traits of mood disorders and depression. The final output from the NAc and regulation of behavioral outcomes is determined by the ChAT activity. This is accomplished by integrating the large number of inputs from various regions, and different neurotransmitter systems that allow fine coordination of ChATs tonic activity. For example, silencing the activity of NAc ChATs increases activity of medium spiny neurons, the main output cells of the NAc ^25^. Based on these data functional implications of the NAc ChATs circuit mapped in this study and its role in depressive behavior most likely depends on the projections unique to NAc, namely, medial, and ventral orbital prefrontal cortex areas, ventral hippocampus and central amygdaloid nucleus. Circuits between these areas and the NAc are known for the role in wide range of behaviors, including depressive behaviors. The complex regulation of the NAc output governed by the groups of neurons within the orbitofrontal cortex and their extensive circuitry with other brain regions has been described ^13, 40, 55^. For example, the PFC-NAC circuitry provides an executive control by mediating task-switching behaviors and inhibiting motor responses ^13^. An increase in the synchronized activity between vHIP and NAc predicted vulnerability to stress in naïve mice and was a marker for mice susceptible to social stress ^40^. In this study we reported several notable findings regarding the connectivity of this region with NAc. We found that layer 2/3 pyramidal cells project directly to the NAc ChATs. This is similar to the wolframin- and Ntf3-expressing layer 2/3 pyramidal neurons, that project to the dorsal striatum, and regulate stress-induced depressive-like behavior in mice ^38^. Furthermore, large body of work have showed that layer 2/3 pyramidal neurons are mostly involved in intracortical circuits ^56, 57^. Our data and data from others ^11, 38^ that found projections from layer 2/3 to subcortical regions. This adds another layer to the cortical circuitry complexity, as the prefrontal cortex is to our knowledge the only area where layer 2/3 pyramidal neurons project outside of the cortex. Also notable is the heterogeneity between layer 2/3 pyramidal cells, since wolframin and Ntf3 expressing neurons do not project to the NAc, while we found neurons in the same region and layer specificity that synapse on the NAc ChATs. Furthermore, it was reported that direct inputs to CPU ChATs arise primarily from the motor areas, and much less from the PFC areas, such as lateral orbital, prelimbic, and cingulate ^21, 22^. This may also imply that prefrontal cortical areas project more heavily to the NAc ChAT cells then hippocampal neurons or amygdala, as suggested before ^7^. Our data support this hypothesis, and further show here that the ventral hippocampus and the medial and ventral orbital cortices project directly and specifically to ventral, but not dorsal ChAT cells. These results point to the complex circuitry between medial orbital cortex and striatum involved in depressive-like behaviors in mice.

However, it is also important to consider that the regulation of depressive phenotype in ChAT-p11 cKO depends mainly on the glutamatergic inputs from the ventral hippocampus, as they were shown here to be significantly reduced in these mice. The influence of distinct glutamatergic projections to the NAc in the regulation of addiction, anxiety, reward, and depressive behavior has been reported previously ^7, 8, 10, 58^ although the specific mechanism reported differs between studies. Britt et al. suggested that ChATs and possibly other interneurons in NAc do not receive direct input from the ventral hippocampus, but indirectly through connections with the medium spiny neurons ^7^. On the other hand, direct inputs from vHIP to Nac were found by others ^10^, and confirmed in this study. LeGates et al. uncovered that strength of the ventral hippocampal input to D1 MSN neurons weakens in chronic stress, and this is linked to changes in reward response ^10^. The role of this circuitry in depressive-like phenotype was further supported by another report showing that glutamatergic transmission from vHIP to NAc MSNs is increased in mice susceptible to stress ^8^. Moreover, this effect is specific to vHIP-NAc circuitry since the stimulation of either mPFC or AMY afferents to the NAc led to resilience. Our study shows for the first-time cell-specific direct glutamatergic input from the ventral hippocampus to the NAc ChATs. Furthermore, this glutamatergic projection appears to have a decisive role in regulating the activity of the NAc ChAT’s in the ChAT-p11 cKO mice, as we found the significant reduction in the number of projections from this region, while projections from PFC, AMY and DR are not changed significantly. It has been proposed that the behavioral output of the NAc depends on the amount of glutamate released from the inputs coming from the PFC and AMY ^7^. It is possible that decreased number of inputs from the vHIP to NAc ChATs disrupts the balance of signals received from other major glutamatergic inputs, including prefrontal cortical areas, central amygdaloid nucleus and parafascicular thalamic nucleus. This would suggest that glutamatergic input disbalance is a key circuitry mechanism of depressive-like phenotype in ChAT-p11 cKO mice (Figs. 5 and 6).

**Figure 6.**
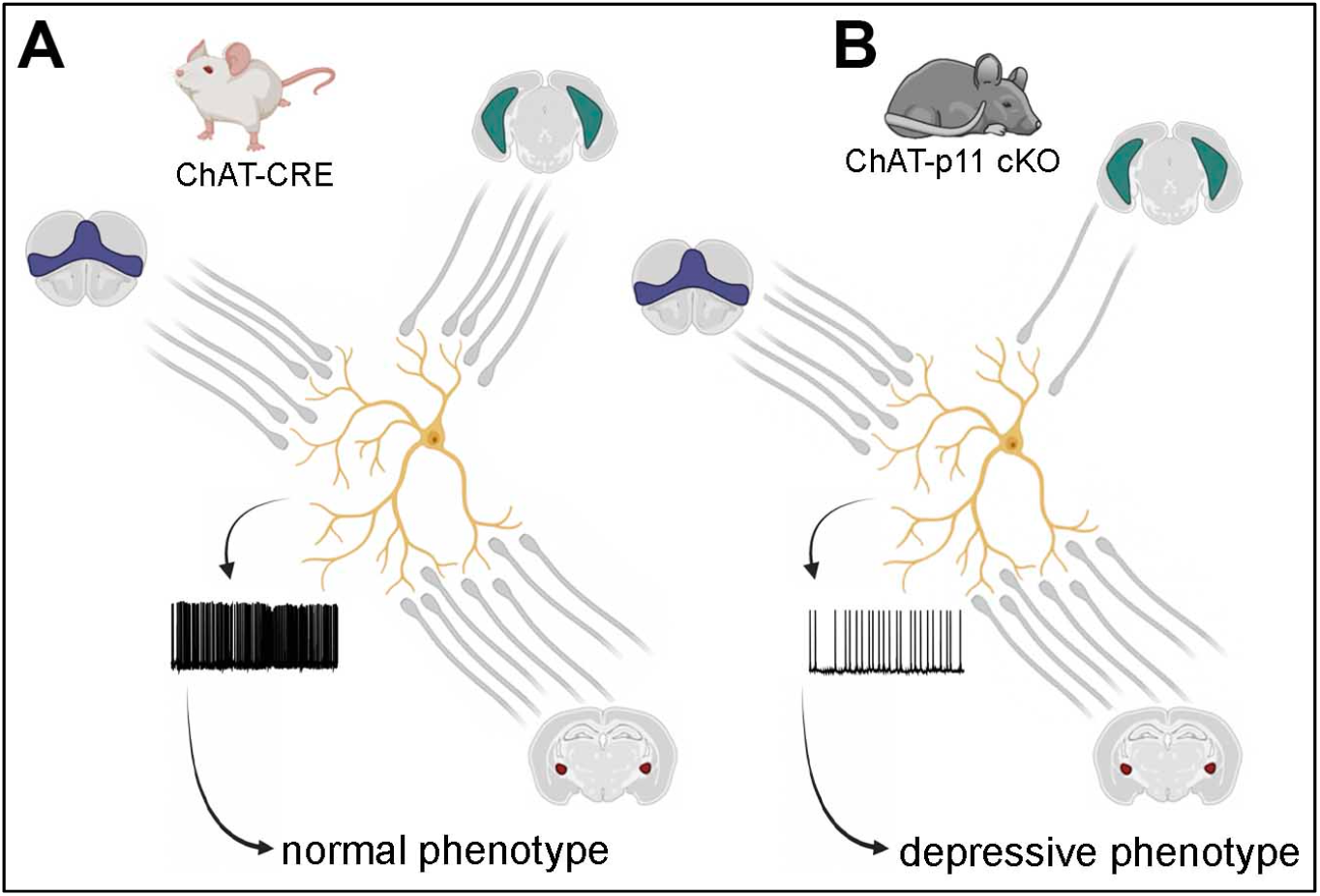
Lack of the inputs from the ventral hippocampus to the nucleus accumbens ChATs represent major circuitry change in the ChAT-p11 cKO mouse model of MDD. **A)** In the healthy mice accumbal ChATs receive glutamatergic projections from three areas, medial orbital PFC (purple), vHIP (teal), and central amygdaloid nucleus (maroon), that appear to be important for their tonic activity and non-depressive phenotype. **B)** In the mouse model of depression inputs from the ventral hippocampus are significantly reduced, tonic activity of the ChATs is severely changed, resulting in a depressive phenotype.

One fascinating outcome outside of the scope of this study is that p11 ablation in ChATs affects specifically the ChAT-vHIP circuitry, but not others examined in this study. One possibility is that p11 affects the development of ChATs in a way that prevents formation of inputs from the vHIP. Genetic ablation impact on ChAT circuitry development has been shown before, where the TrkA ablation impairs the basal forebrain-vHIP ChAT circuitry, specifically its laminar pattern and the number of projections ^59^. In rodents ChATs development is tightly regulated by the NGF and BDNF signaling cascades trough their receptors, p75, TrkA, and TrkB, where the NGF/p75 determines the number of striatal ChATs ^59, 60^, while the BDNF/TrkB regulate growth and complexity of neurons, including loss of dendritic spines, and diminished nigral-striatal projections ^61^. The mechanism by which p11 affects ChAT development may rely on a complex cascade involving plasmin, BDNF, NGF, and Anxa2, an important binding partner of p11. Results from the *in vitro* studies support this hypothesis. For example, it has been shown that NGF-induced neuritogenesis depends on the Anxa2-mediated plasmin generation ^62^, while p11 has been shown to be necessary for the BDNF-mediated effect on dendritic length and spine density ^63^. However, the exact mechanism of p11-induced changes in NAc ChAT circuitry shown here remain to be elucidated.

### Limitations of the study related to the methodology

The difficulty of labeling sparse populations and areas deep within the brain, such as NAc, has so far prevented mapping of the interneuronal subtypes in NAc. The optimal infection of scant populations, such as ChATs, requires larger number of viral particles and higher volume for injection, which then brings a risk of off target infections in adjacent areas with the same type of the cell. For example, olfactory bulb granule cells as well as olfactory tubercle neurons were very often labeled due to the infection of olfactory tubercle neurons and piriform cortex. To circumvent this constraint various volumes of rabies viral particles were injected. While labeled cells were detected in the same areas of the brain, much smaller number of cells was observed with the smaller volume. Thus, smaller volumes ensured the proper regional quantification, but the optimal visualization of the projection sites in the whole brain tissue had to be performed with the higher volume. Furthermore, in this study even the smaller volume of the virus injected prevented us to discern between the projections to the core and shell of the NAc, as the virus infected both areas of the NAc. We also detected some areas, particularly in the regions further away from the injection site, such as locus coeruleus or medulla, with the number of projections in single digits. This implied that the number of viral particles defined by the titer of the virus and the volume injected, is very important for its propagation and consequently a number of labeled cells and quality of labeling, as suggested previously ^20^. Therefore, while this study used the approach in experimental design and rigorous analysis of virally labeled projection neurons, previously design by others ^21, 22, 42^, these circuitries will need further confirmation by other methods, such as electrophysiology.

***In summary***, given the key role of the ChATs, and especially NAc ChATs in mood disorders, we elucidated the connectivity of the accumbal ChATs on the level of the whole brain and examined this connectivity in the context of the mouse model of depression. Specific behavioral patterns and clinical symptoms of depression depend on the correlations between specific networks between multiple brain areas, their interactions, and cell-specific circuits between these regions. In this context, functional significance of our data warrants further mechanistic studies that will elucidate the role of NAc ChAT inputs from various brain regions and how they govern reward behavior and dysfunction in mood disorders. Furthermore, this study may further our understanding of the functional changes in the patients suffering from neurodegenerative diseases, such as Alzheimer’s, Parkinson’s and Huntington’s which have disrupted acetylcholine signaling in the striatum, and often exhibit psychiatric symptoms.

## Funding and acknowledgements

This work was supported by the Fisher Foundation (to PG), RF1AG059770 and NIDA P30 DA035756 Pilot Project Award (to A Milosevic).

We thank Dr. Nathaniel Heintz for a generous gift of the ChAT-CRE and ChAT-TRAP mice lines (project Gene Expression Nervous System Atlas (GENSAT) Project, NINDS Contracts N01NS02331 & HHSN271200723701C to The Rockefeller University). We are grateful to Dr. Urlich Muller (John Hopkins School of Medicine, Baltimore, MD) for the Cux2-CRE mouse line. We thank Dr. Katia Manova-Todorova, the head of MSKCC Molecular Imaging Core Facility, Dr. Jerry Chang, and Dr. Thomas Liebman for help with the imaging. The authors thank Elizabeth Griggs for expert help with graphics. Special thanks to Drs. Inez Ibanez-Tallon, Luca Parolari and Jennifer Warner-Schmidt for helpful suggestions during this study. We thank Dr. Marc Flajolet who read the manuscript and offered thoughtful suggestions. We extend special thanks to Dina Becaj, a participant of the Rockefeller University Summer Science Research Program, who helped with the cell quantification experiments.

Role of Authors: all authors take responsibility for the accuracy of the data and analysis. All authors had full access to all the data in the study. Study concept and design: AM, LM, PG. Acquisition of the data: AM, LM, MK, TF, PDCV, YR, SF. Analysis and interpretation of the data: AM, LM, MK, YR, SF. Drafting of the paper: AM, LM. Obtained funding: AM, PG. Technical support: MK, TF, PCDV. Study supervisor: AM.

## Conflict of interest

The authors declare no competing interests.

## Methods

### Animals and reagents

Adult Chat-CRE (GM60) mice obtained from GENSAT (www.gensat.org) and Chat+ cell specific p11 knockout mice Chat-CRE/p11 floxed constitutive were used in this study. Cell-type specific p11 knockout mouse line was generated by breeding the p11 floxed mice to Chat-CRE mice, as described previously ^32^. In addition, this mouse line was also breed to ChAT-TRAP line ^33^. We utilized p11 knockout mouse as a model of depression ^30^. The mouse lines were maintained in a C57BL/6 background. Additionally, Cux2-CRE line was used (kindly donated by Ulrich Muller, Johns Hopkins School of Medicine, Baltimore, MD). Camk2a-CRE line was obtained from the Jackson laboratory (B6.Cg-Tg(Camk2a-cre)T29-1Stl/J). Mice were kept at the 12 hours light/dark cycle and water and food ad libitum at the Rockefeller University animal facility. Mice of both sexes were selected for experiments when they were 8 to 16 weeks old. For all experiments the age and sex matched mice were used as controls. All procedures were approved by the institutional IACUC and adhere to the ARRIVE guidelines.

### Viruses

Adenoviral vectors AAV2.CMV.PI.Cre.rBG and AAV2.CMV.HI.GFP-Cre.SV40 adenoviral vectors ^19^ were purchased from University of North Carolina at Chapel Hill Vector Core. Rabies virus vectors EnvA G-deleted Rabies-GFP or EnvA G-deleted Rabies-mCherry ^17^ were purchased from GT3 Core Facility at Salk Institute.

### Stereotaxic injections

We performed three types of injection: helper virus was injected into NAC, followed by the injection of the rabies virus at the different angle to avoid infection and replication of the virus in the dorsal striatum. Stereotaxic surgeries were performed under general ketamine-xylazine anesthesia. Mice were placed in the stereotaxic apparatus and injection was guided by the AngleTwo program. Coordinates for the nucleus accumbens injection were +1.10 mm mediolateral, +1.80 mm anteroposterior, and -4.60 mm dorsoventral from bregma. Coordinates for DSTR injections were: +1.80mm lateral, 0.70 anterior and -3.50mm dorsoventral from the bregma. Each mouse was injected with 0.3-1mL of the viral suspension using the 10 mL Hamilton syringe and 33g Hamilton needle over 10 minutes and an infusion pump (World Precision Instruments). Two consecutive injections were performed on each mouse. The helper virus mixture, consisting of the equal volume of AAV2.CMV.PI.Cre.rBG and AAV2.CMV.HI.GFP-Cre.SV40 adenoviral vectors, was injected at the volume of 0.8 µl, followed by the 0.3 µl of the EnvA G-deleted Rabies-GFP or EnvA G-deleted Rabies-mCherry 4-5 weeks after the first injection. The titer of the viruses was 0.9-1cfu ×10^6^. For the injections into NAC, the second injections were performed at the 25° angle to avoid infecting the cells in the CPU with the rabies viral vector. Mice were euthanized and analyzed one week after the rabies virus injection. Since this technique involves stereotaxic injection into the ventral part of the striatum, there is a possibility of unintended infection of the ChAT cells in the dorsal striatum along the injection route. Thus, two sets of injections were done: one into NAc and one into dorsal striatum right above the NAc injection with the smaller (0.3 µl) and larger (0.8 µl) volume of the rabies virus. The analysis with the larger volume was used for the scanning of the whole brain for the representative images, because it enables easier visualization of the projection sites.

### Tissue processing and immunocytochemistry

One week after the injection of the EnvA G-deleted Rabies viral vector animals were deeply anesthetized with Nembutal and intracardially perfused with 4% paraformaldehyde solution and cryopreserved through series of sucrose dilutions. Brains were cryopreserved in a series of sucrose solutions in 0.01M PBS and embedded in the embedding media (Tissue-Tek). Series of 30 microns-thick coronal sections throughout the whole brain were sectioned on the cryostat. Immunocytochemistry was performed as described before ^34^. In short, cryostat sections were incubated with the solution of 5% normal goat serum in PBS for 1 hr, followed by overnight incubation with the primary antibodies and appropriate Alexa Fluor (Invitrogen) secondary antibodies. Primary antibodies used were anti-GFP (1/500, raised in chicken, Abcam) and mCherry (1/500, raised in rabbit, Abcam). Sections were coverslipped with ProLong mounting solution.

### Imaging and quantification

Imaging was performed in two ways. For representative images of the brain areas, tissue sections were first labeled with the anti-GFP or anti-mCherry antibodies, depending on the type of the RV injected. Then, images were selected based on the quality of expression and imaged at the confocal microscope (LSM710, Zeiss). Images shown throughout the paper were processed in ImageJ (for the scale bars) and Photoshop. In Photoshop, regions with labeled cells were enlarged to enable better visual representation of projecting neurons. Images were modified only with the brightness and contrast Photoshop’s image modification tool. Schematic diagrams were created with Biorender.com.

For quantification, three biological and technical replicate samples were used for control and ChAT-p11 cKO. Imaging for quantification was performed on a series of sections that contained every other 30 micron-thick section throughout the whole brain. Imaging was performed with the Mirax Scan scanning system. The resulting low magnification images represented 5×5 tiled images of a whole brain section. These images were loaded into the PanoramicView software, and areas with GFP-labeled cells were outlined in their respective brain regions. The expression sites delineations were made in accordance with The Mouse Brain atlas ^35^ and Scalable Brain Atlas (http://scalablebrainatlas.incf.org/) as reference. MetaMorph Image Analysis software was used for the automatic quantification of GFP- or mCherry-labeled cell bodies in each region, and ImageJ was used for the semi-automatic conformation of the cell body numbers. Differential enrichment of cell bodies between different regions and respective graph bar chart was completed using the Prism software. For this analysis only areas that appear in at least two samples were analyzed. Areas at the injection sites, as well as septal, basal forebrain, extended amygdala, and cortical-amygdala transitory areas that contain large number of ChAT cells were not included in the analysis. We used Paxinos atlas to outline the brain regions, and the abbreviations for these regions are shown in Table 1.

**Table 1.**
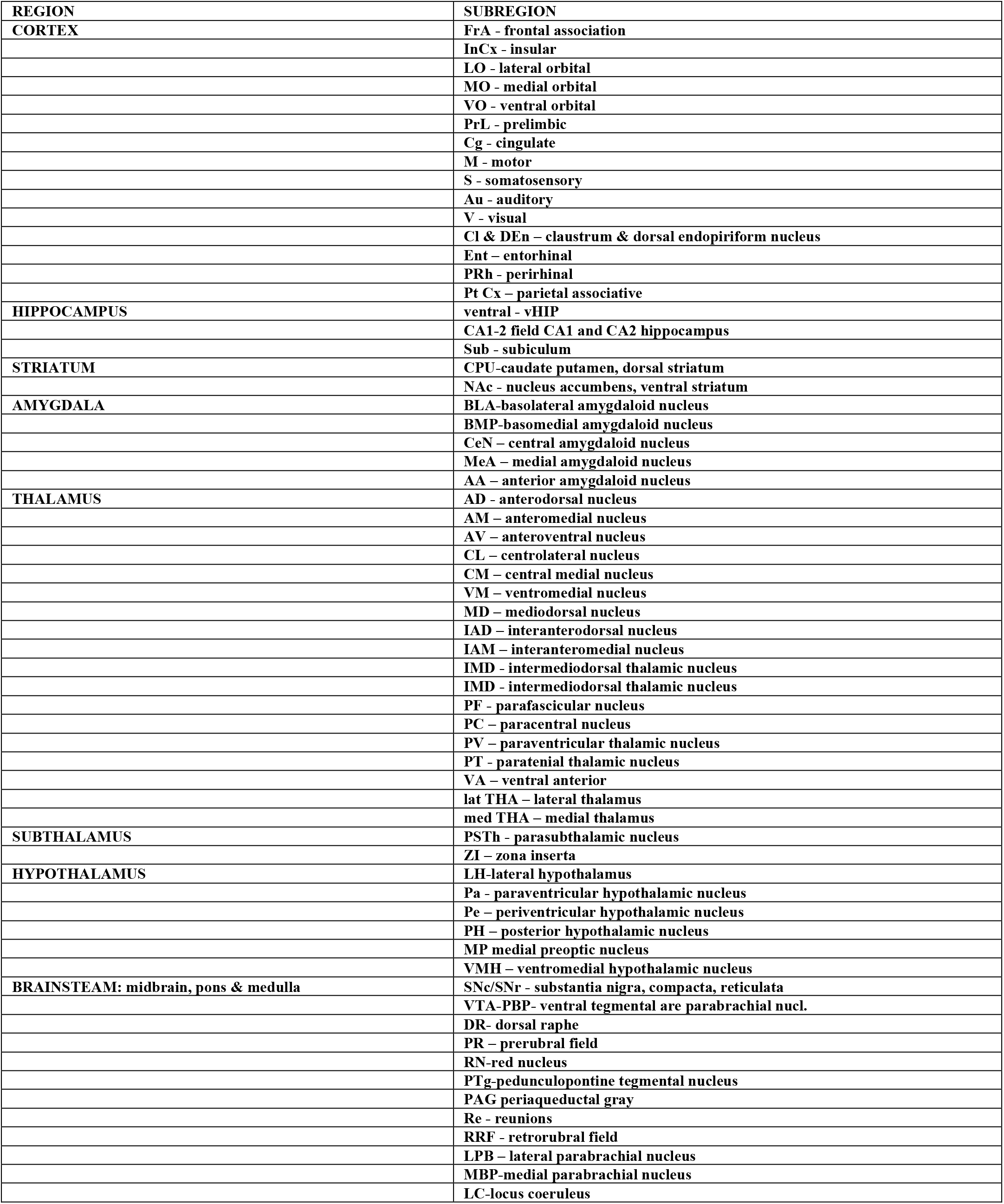
List of anatomical structure abbreviations used in the study.

### Optogenetics

Mice were anesthetized with a ketamine/xylazine cocktail and underwent stereotaxic surgery to inject serotype 5 adeno-associated viruses (AAV) encoding CaMK2a-ChR2 (H134R) –eYFP (UNC Viral Vector Core, Chapel Hill, North Carolina). The virus was injected bilaterally into the ventral hippocampus (from bregma: anterior/posterior: –3.8, lateral: +3.0, dorsal/ventral: –4.5 from top of skull) at a rate of 0.1 ml per minute for 10 minutes. Mice recovered for 5-6 weeks before being subjected to electrophysiological experiments. For electrophysiological recordings, field light stimulation of ChR2-expressing vHIP terminals in NAc neurons was done through a 40x objective using a SPECTRA X LED light engine (Lumencor, OR, USA).

### Electrophysiology

Mice between 8 and 12 weeks of age were euthanized with CO2. Coronal slices (300 μm thickness) were cut using a Vibratome 1000 Plus (Leica Microsystems, USA) at 2 °C in a NMDG-containing cutting solution (in mM): 105 NMDG (N-Methyl-D-glucamine), 105 HCl, 2.5 KCl, 1.2 NaH2PO4, 26 NaHCO3, 25 Glucose, 10 MgSO4, 0.5 CaCl2, 5 L-Ascorbic Acid, 3 Sodium Pyruvate, 2 Thiourea (pH was around 7.4, with osmolarity of 295–305 mOsm). After cutting, slices recovered for 15 minutes in the same cutting solution at 35 °C and for 1 h at room temperature (RT) in aCSF recording solution containing (in mM): 125 NaCl, 25 NaHCO3, 2.5 KCl, 1.25 NaH2PO4, 2 CaCl2, 1 MgCl2 and 25 glucose (bubbled with 95% O2 and 5% CO2) (see below). Whole-cell patch-clamp recordings were performed with a Multiclamp 700B/Digidata1550A system (Molecular Devices, Sunnyvale CA, USA), an upright Olympus BX51WI microscope equipped with the appropriate filters (Olympus, Japan) and a SPECTRA X LED light engine (Lumencor, OR, USA). The slice was placed in a recording chamber (RC-27L, Warner Instruments, USA) and constantly perfused with oxygenated aCSF at 24 °C (TC-324B, Warner Instruments, USA) at a rate of 1.5–2.0 ml/min. The intracellular solution contained (in mM): 126 K-gluconate, 4 NaCl, 1 MgSO4, 0.02 CaCl2, 0.1 BAPTA, 15 glucose, 5 HEPES, 3 ATP, 0.1 GTP (pH 7.3). We used whole-cell current-clamp for electrophysiology recordings in layer II/III PFC neurons. For measuring the membrane potential 30 seconds of recording were binned into 0.5 ms bins and fitted with a Gaussian. For measuring the action potential firing, small currents were injected to the cells to bring the membrane potential at -70 mV. Consecutive 1 s current steps of 50 pA starting from -100 pA were injected to induce depolarization. Action potential threshold was measured from the first action potential to avoid any confounding effects of adaptation. For electrophysiology recording in nucleus accumbens we used whole-cell current-clamp to record the tonic activity of ChAT neurons. Consecutively, we used whole-cell voltage-clamp configuration to record evoked glutamatergic responses in ChAT or MSN neurons. For this, we used field light stimulation of ChR2-expressing vHIP terminals in NAc in the presence of 30 µM bicuculine (to block GABAergic transmission).

## Notes

### Competing Interest Statement

The authors have declared no competing interest.

## References

1. Zahm DS, Brog JS. On the significance of subterritories in the “accumbens” part of the rat ventral striatum. Neuroscience 1992; 50(4): 751–767.

2. O’Donnell P, Lavin A, Enquist LW, Grace AA, Card JP. Interconnected parallel circuits between rat nucleus accumbens and thalamus revealed by retrograde transynaptic transport of pseudorabies virus. J Neurosci 1997; 17(6): 2143–2167.

3. Meredith GE, Baldo BA, Andrezjewski ME, Kelley AE. The structural basis for mapping behavior onto the ventral striatum and its subdivisions. Brain Struct Funct 2008; 213(1-2): 17–27.

4. Zahm DS. An integrative neuroanatomical perspective on some subcortical substrates of adaptive responding with emphasis on the nucleus accumbens. Neurosci Biobehav Rev 2000; 24(1): 85–105.

5. Russo SJ, Nestler EJ. The brain reward circuitry in mood disorders. Nat Rev Neurosci 2013; 14(9): 609–625.

6. Price JL, Drevets WC. Neurocircuitry of mood disorders. Neuropsychopharmacology : official publication of the American College of Neuropsychopharmacology 2010; 35(1): 192–216.

7. Britt JP, Benaliouad F, McDevitt RA, Stuber GD, Wise RA, Bonci A. Synaptic and behavioral profile of multiple glutamatergic inputs to the nucleus accumbens. Neuron 2012; 76(4): 790–803.

8. Bagot RC, Parise EM, Peña CJ, Zhang HX, Maze I, Chaudhury D et al. Ventral hippocampal afferents to the nucleus accumbens regulate susceptibility to depression. Nat Commun 2015; 6: 7062.

9. Christoffel DJ, Golden SA, Walsh JJ, Guise KG, Heshmati M, Friedman AK et al. Excitatory transmission at thalamo-striatal synapses mediates susceptibility to social stress. Nat Neurosci 2015; 18(7): 962–964.

10. LeGates TA, Kvarta MD, Tooley JR, Francis TC, Lobo MK, Creed MC et al. Reward behaviour is regulated by the strength of hippocampus-nucleus accumbens synapses. Nature 2018; 564(7735): 258–262.

11. Thompson RH, Swanson LW. Hypothesis-driven structural connectivity analysis supports network over hierarchical model of brain architecture. Proc Natl Acad Sci U S A 2010; 107(34): 15235–15239.

12. Smith Y, Raju DV, Pare JF, Sidibe M. The thalamostriatal system: a highly specific network of the basal ganglia circuitry. Trends Neurosci 2004; 27(9): 520–527.

13. Sesack SR, Grace AA. Cortico-Basal Ganglia reward network: microcircuitry. Neuropsychopharmacology 2010; 35(1): 27–47.

14. Warner-Schmidt JL, Schmidt EF, Marshall JJ, Rubin AJ, Arango-Lievano M, Kaplitt MG et al. Cholinergic interneurons in the nucleus accumbens regulate depression-like behavior. Proc Natl Acad Sci U S A 2012; 109(28): 11360–11365.

15. Gerfen CR, Sawchenko PE. An anterograde neuroanatomical tracing method that shows the detailed morphology of neurons, their axons and terminals: immunohistochemical localization of an axonally transported plant lectin, Phaseolus vulgaris leucoagglutinin (PHA-L). Brain Res 1984; 290(2): 219–238.

16. Bolam JP, Hanley JJ, Booth PA, Bevan MD. Synaptic organisation of the basal ganglia. Journal of anatomy 2000; 196 (Pt 4): 527–542.

17. Wall NR, Wickersham IR, Cetin A, De La Parra M, Callaway EM. Monosynaptic circuit tracing in vivo through Cre-dependent targeting and complementation of modified rabies virus. Proc Natl Acad Sci U S A 2010; 107(50): 21848–21853.

18. Wall NR, Wickersham IR, Cetin A, De La Parra M, Callaway EM. Monosynaptic circuit tracing in vivo through Cre-dependent targeting and complementation of modified rabies virus. Proc Natl Acad Sci U S A 2010; 107(50): 21848–21853.

19. Watabe-Uchida M, Zhu L, Ogawa SK, Vamanrao A, Uchida N. Whole-brain mapping of direct inputs to midbrain dopamine neurons. Neuron 2012; 74(5): 858–873.

20. Wall NR, De La Parra M, Callaway EM, Kreitzer AC. Differential innervation of direct-and indirect-pathway striatal projection neurons. Neuron 2013; 79(2): 347–360.

21. Guo Q, Wang D, He X, Feng Q, Lin R, Xu F et al. Whole-brain mapping of inputs to projection neurons and cholinergic interneurons in the dorsal striatum. PLoS One 2015; 10(4): e0123381.

22. Klug JR, Engelhardt MD, Cadman CN, Li H, Smith JB, Ayala S et al. Differential inputs to striatal cholinergic and parvalbumin interneurons imply functional distinctions. Elife 2018; 7.

23. Rymar VV, Sasseville R, Luk KC, Sadikot AF. Neurogenesis and stereological morphometry of calretinin-immunoreactive GABAergic interneurons of the neostriatum. J Comp Neurol 2004; 469(3): 325–339.

24. Virk MS, Sagi Y, Medrihan L, Leung J, Kaplitt MG, Greengard P. Opposing roles for serotonin in cholinergic neurons of the ventral and dorsal striatum. Proc Natl Acad Sci U S A 2016; 113(3): 734–739.

25. Witten IB, Lin SC, Brodsky M, Prakash R, Diester I, Anikeeva P et al. Cholinergic interneurons control local circuit activity and cocaine conditioning. Science 2010; 330(6011): 1677–1681.

26. Collins AL, Aitken TJ, Huang IW, Shieh C, Greenfield VY, Monbouquette HG et al. Nucleus Accumbens Cholinergic Interneurons Oppose Cue-Motivated Behavior. Biol Psychiatry 2019; 86(5): 388–396.

27. Alexander B, Warner-Schmidt J, Eriksson T, Tamminga C, Arango-Lievano M, Ghose S et al. Reversal of depressed behaviors in mice by p11 gene therapy in the nucleus accumbens. Sci Transl Med 2010; 2(54): 54ra76.

28. Cheng J, Umschweif G, Leung J, Sagi Y, Greengard P. HCN2 Channels in Cholinergic Interneurons of Nucleus Accumbens Shell Regulate Depressive Behaviors. Neuron 2019; 101(4): 662-672.e665.

29. Warner-Schmidt JL, Schmidt EF, Marshall JJ, Rubin AJ, Arango-Lievano M, Kaplitt MG et al. Cholinergic interneurons in the nucleus accumbens regulate depression-like behavior. Proc Natl Acad Sci U S A 2012; 109(28): 11360–11365.

30. Svenningsson P, Chergui K, Rachleff I, Flajolet M, Zhang X, El Yacoubi M et al. Alterations in 5-HT1B receptor function by p11 in depression-like states. Science 2006; 311(5757): 77–80.

31. Zhang L, Su TP, Choi K, Maree W, Li CT, Chung MY et al. P11 (S100A10) as a potential biomarker of psychiatric patients at risk of suicide. J Psychiatr Res 2011; 45(4): 435–441.

32. Warner-Schmidt JL, Vanover KE, Chen EY, Marshall JJ, Greengard P. Antidepressant effects of selective serotonin reuptake inhibitors (SSRIs) are attenuated by antiinflammatory drugs in mice and humans. Proc Natl Acad Sci U S A 2011; 108(22): 9262–9267.

33. Doyle JP, Dougherty JD, Heiman M, Schmidt EF, Stevens TR, Ma G et al. Application of a translational profiling approach for the comparative analysis of CNS cell types. Cell 2008; 135(4): 749–762.

34. Milosevic A, Liebmann T, Knudsen M, Schintu N, Svenningsson P, Greengard P. Cell-and region-specific expression of depression-related protein p11 (S100a10) in the brain. J Comp Neurol 2017; 525(4): 955–975.

35. Franklin KBJ, Paxinos G. The Mouse Brain in Stereotaxic Coordinates, Compact Third Edition. Academic Press-Elsevier: New York, 2008.

36. Schmidt EF, Warner-Schmidt JL, Otopalik BG, Pickett SB, Greengard P, Heintz N. Identification of the cortical neurons that mediate antidepressant responses. Cell 2012; 149(5): 1152–1163.

37. Franco SJ, Gil-Sanz C, Martinez-Garay I, Espinosa A, Harkins-Perry SR, Ramos C et al. Fate-restricted neural progenitors in the mammalian cerebral cortex. Science 2012; 337(6095): 746–749.

38. Shrestha P, Mousa A, Heintz N. Layer 2/3 pyramidal cells in the medial prefrontal cortex moderate stress induced depressive behaviors. Elife 2015; 4.

39. Stuber GD, Sparta DR, Stamatakis AM, van Leeuwen WA, Hardjoprajitno JE, Cho S et al. Excitatory transmission from the amygdala to nucleus accumbens facilitates reward seeking. Nature 2011; 475(7356): 377–380.

40. Hultman R, Ulrich K, Sachs BD, Blount C, Carlson DE, Ndubuizu N et al. Brain-wide Electrical Spatiotemporal Dynamics Encode Depression Vulnerability. Cell 2018; 173(1): 166-180.e114.

41. Chaudhury D, Liu H, Han MH. Neuronal correlates of depression. Cell Mol Life Sci 2015.

42. Haubensak W, Kunwar PS, Cai H, Ciocchi S, Wall NR, Ponnusamy R et al. Genetic dissection of an amygdala microcircuit that gates conditioned fear. Nature 2010; 468(7321): 270–276.

43. Tye KM, Prakash R, Kim SY, Fenno LE, Grosenick L, Zarabi H et al. Amygdala circuitry mediating reversible and bidirectional control of anxiety. Nature 2011; 471(7338): 358–362.

44. Paré D. Role of the basolateral amygdala in memory consolidation. Prog Neurobiol 2003; 70(5): 409–420.

45. Ventura-Silva AP, Melo A, Ferreira AC, Carvalho MM, Campos FL, Sousa N et al. Excitotoxic lesions in the central nucleus of the amygdala attenuate stress-induced anxiety behavior. Front Behav Neurosci 2013; 7: 32.

46. Niu JG, Yokota S, Tsumori T, Oka T, Yasui Y. Projections from the anterior basomedial and anterior cortical amygdaloid nuclei to melanin-concentrating hormone-containing neurons in the lateral hypothalamus of the rat. Brain Res 2012; 1479: 31–43.

47. Pérez CA, Stanley SA, Wysocki RW, Havranova J, Ahrens-Nicklas R, Onyimba F et al. Molecular annotation of integrative feeding neural circuits. Cell Metab 2011; 13(2): 222–232.

48. Stanley S, Pinto S, Segal J, Pérez CA, Viale A, DeFalco J et al. Identification of neuronal subpopulations that project from hypothalamus to both liver and adipose tissue polysynaptically. Proc Natl Acad Sci U S A 2010; 107(15): 7024–7029.

49. Brown MT, Tan KR, O’Connor EC, Nikonenko I, Muller D, Lüscher C. Ventral tegmental area GABA projections pause accumbal cholinergic interneurons to enhance associative learning. Nature 2012; 492(7429): 452–456.

50. Chaudhury D, Walsh JJ, Friedman AK, Juarez B, Ku SM, Koo JW et al. Rapid regulation of depression-related behaviours by control of midbrain dopamine neurons. Nature 2013; 493(7433): 532–536.

51. Waselus M, Galvez JP, Valentino RJ, Van Bockstaele EJ. Differential projections of dorsal raphe nucleus neurons to the lateral septum and striatum. J Chem Neuroanat 2006; 31(4): 233–242.

52. Cachope R, Mateo Y, Mathur BN, Irving J, Wang HL, Morales M et al. Selective activation of cholinergic interneurons enhances accumbal phasic dopamine release: setting the tone for reward processing. Cell Rep 2012; 2(1): 33–41.

53. Berton O, McClung CA, Dileone RJ, Krishnan V, Renthal W, Russo SJ et al. Essential role of BDNF in the mesolimbic dopamine pathway in social defeat stress. Science 2006; 311(5762): 864–868.

54. Epstein J, Pan H, Kocsis JH, Yang Y, Butler T, Chusid J et al. Lack of ventral striatal response to positive stimuli in depressed versus normal subjects. Am J Psychiatry 2006; 163(10): 1784–1790.

55. Pinto A, Sesack SR. Limited collateralization of neurons in the rat prefrontal cortex that project to the nucleus accumbens. Neuroscience 2000; 97(4): 635–642.

56. Yoshimura Y, Dantzker JL, Callaway EM. Excitatory cortical neurons form fine-scale functional networks. Nature 2005; 433(7028): 868–873.

57. Shepherd GM. Corticostriatal connectivity and its role in disease. Nat Rev Neurosci 2013; 14(4): 278–291.

58. Muir J, Tse YC, Iyer ES, Biris J, Cvetkovska V, Lopez J et al. Ventral Hippocampal Afferents to Nucleus Accumbens Encode Both Latent Vulnerability and Stress-Induced Susceptibility. Biol Psychiatry 2020; 88(11): 843–854.

59. Sanchez-Ortiz E, Yui D, Song D, Li Y, Rubenstein JL, Reichardt LF et al. TrkA gene ablation in basal forebrain results in dysfunction of the cholinergic circuitry. J Neurosci 2012; 32(12): 4065–4079.

60. Ward NL, Hagg T. p75(NGFR) and cholinergic neurons in the developing forebrain: a re-examination. Brain Res Dev Brain Res 1999; 118(1-2): 79-91.

61. Li Y, Yui D, Luikart BW, McKay RM, Li Y, Rubenstein JL et al. Conditional ablation of brain-derived neurotrophic factor-TrkB signaling impairs striatal neuron development. Proc Natl Acad Sci U S A 2012; 109(38): 15491–15496.

62. Jacovina AT, Zhong F, Khazanova E, Lev E, Deora AB, Hajjar KA. Neuritogenesis and the nerve growth factor-induced differentiation of PC-12 cells requires annexin II-mediated plasmin generation. J Biol Chem 2001; 276(52): 49350–49358.

63. Park SW, Nhu lH, Cho HY, Seo MK, Lee CH, Ly NN et al. p11 mediates the BDNF-protective effects in dendritic outgrowth and spine formation in B27-deprived primary hippocampal cells. J Affect Disord 2016; 196: 1-10.

